# Estimating individuals’ exposure to predation risk in group-living baboons, *Papio anubis*

**DOI:** 10.1101/2023.06.06.543865

**Authors:** Alexandre Suire, Itsuki Kunita, Roi Harel, Margaret Crofoot, Mathew Mutinda, Maureen Kamau, James M. Hassel, Suzan Murray, Akiko Matsumoto-Oda

## Abstract

In environments with multiple predators, the vulnerabilities attached to the spatial positions of group-living prey are not uniform and depend on the hunting styles of the predators. Coursing predators, mainly canids and hyenas, follow their prey over long distances and attack open areas, making individuals at the edge of the group more dangerous than those at the center (marginal predation). In contrast, ambush predators, mainly cats, approach their prey undetected and appear randomly anywhere in the group; therefore, isolated individuals are at a greater risk of predation. However, identifying individuals at high risk of predation requires the simultaneous recording of predator locations and direct observation of predation events, which are both difficult. Therefore, several theoretical methods have been proposed to assess predation risk (predation risk proxies). In a group of wild anubis baboons exposed to predation by leopards, lions, and hyenas, we calculated predation risk proxies using movement data collected from global positioning system (GPS) collars and found that adult males were on the edge of the group with a higher risk of predation (Hypothesis 1). In addition, adult males were more isolated within this group (Hypothesis 2). None of the predation risks differed among the other age-sex classes. The most dominant male was expected to be in the safety center of the group (Hypothesis 3) but was isolated on the periphery, like the other males. Therefore, we discussed why adult males were more peripheral and isolated.

## Introduction

Under the threat of predation, animals are often tightly grouped, with all group members benefiting from a reduction in predation risk through several mechanisms, including dilution, many-eyes, and confusion effects [1,2]. However, predation risk is often unequal among individuals with different attributes and states within a group. For example, predators may select prey based on their individual vulnerabilities [3–5]because predators can conserve hunting energy or reduce the risk of injury by selecting prey in poor conditions [6–8]. In many taxa, predators have shown evidence of capturing young, old, sick, weak, injured, or inexperienced individuals from prey populations that are higher than expected [4,9,10].

The spatial position within the prey group has also been predicted to lead to vulnerability to predation [11–13].

Studies on wild primates have contributed to the empirical knowledge that different individuals occupy different positions in a group. This is because, unlike birds, fish, or other mammals, primates form groups large enough for human observation, and individuals have been identified in most studies. In summary, individuals that occupy the center of a group are dominant [14,15], highly sociable [16], stay longer in the group [17], and are female and immature [18,19].

Males are located in the forward direction of the group travelling, where there is high risk of encountering predators [15,20–22]. However, opposite patterns have been reported in other animals including primates, where the predation risk was highest in the center of the group [23,24] or at the rear of the group [21,25,26].

One possible reason for the empirical ambiguity may be related to the data collection methods. In previous studies, the spatial positions of individuals were obtained by recording the order in which animals passed through anchor points, such as roads, or by measuring individual distances from the anchor using foot observers. Recently, however, human observers have been found to affect primates with varying degrees of habituation as well as predators [27]. Specifically, shy individuals tend to avoid human observers, creating a bias in the spatial positioning data. Hunted individuals may have been observed under these conditions. Global Positioning Systems (GPS) can effectively avoid this problem by recording the positions of individuals even in the absence of human observers.

Alternatively, another reason could be related to the hunting styles of predators. Predator-hunting styles are generally divided into coursing and ambushing. Because coursing predators (primarily canids and hyenas) chase their prey over long distances and attack from open areas, individuals at the edge of a group are more likely to encounter predators than individuals at the center and are more dangerous (‘ marginal predation’ : [12,28]). In contrast, ambush predators (mainly felines) approach their prey unnoticed and randomly appear anywhere in the group; therefore, the predation risk is higher for isolated individuals [29–32]. This indicates that individuals vulnerable to one type of predator may not be as vulnerable to another type of predator. Most studies on predator-prey responses have focused on only one predator species [33,34], despite multiple predator species in an ecosystem. Studies that account for the risk of multiple predators are required to derive a general rule regarding which positions are vulnerable to predation.

In this study, we used multiple GPS devices simultaneously on a group of individually identified wild baboons to re-examine the age, sex, and dominance of individuals in positions expected to be vulnerable to each of the two types of terrestrial predators, under conditions that minimized the influence of human observers. Although the sample size of predation events was small, a previous study reported that predators were more likely to hunt adults than juveniles and males than females [35]. Instead of directly observing predation events, we calculated individual predation risk using five alternative indices (proxies) that theoretical studies have proposed to predict the predation risk faced by individual prey in two-dimensional space. We tested the following three hypotheses. Hypothesis 1 was that adult males are located peripherally, where coursing predation is most likely to occur. Hypothesis 2 states that the isolated individuals vulnerable to ambush predators are adult males. Hypothesis 3 states that dominant individuals, especially the alpha male, occupy the center of the group.

## Methods

### Ethical statement

The research activities described in this paper were approved by the National Commission for Science, Technology, and Innovation of Kenya (Permit No. NACOSTI/P/16/84320/12475), the Kenya Wildlife Service (KWS/BRP/5001), and UC Davis IACUC Protocol (D16-00272:20442).

### Study site and subjects

This study was conducted at the Mpala Research Centre (0°20’ N, 36°50’ E) in Laikipia Plateau, Kenya. The surrounding environment included bushes and riverine woodlands dominated by *Acacia* spp. The annual rainfall at Mpala is approximately 700 mm, with trilateral peaks in the order of April–May, October–November, and July–August [36].

However, rainfall was highly variable each year, and in July 2019, only approximately half (37 mm) of the normal rainfall (56 mm, 1998–2018) fell, providing observers with good visibility.

The wild anubis baboon (*Papio anubis*) ‘ AI’ group has been studied since 2011. Between July and August 2019, 63 individuals including infants were included in this group. Each day, we recorded individual survival and female estrus status in the morning and evening at the sleeping sites. The number of estrous females during the analysis period was 1.80 ± 1.70, which was not different from the 1.62 ± 0.91 for the rest of the study period (t-test, t=-0.30, n1=5, n2=34, p=0.77). During the study period, excluding the first 5 days, we recorded behaviors by ad libitum observation, and the alpha male was determined by direct behavioral observations, such as attacks or displacements of other individuals in the group. The alpha male of this study period, *MG*, was assumed to have been born in this group because his body size was a late juvenile-early adolescent when he was identified. Leopards, lions, and hyenas were sympatric with the AI group. The predation rate on the AI group from 2011 to 2016 was estimated to be 0.06 individuals/year [37].

### Collaring methods and GPS recorded data

Baboons were captured using walk-in traps (1.8 m^3^) baited with maize and placed near the sleeping sites of the groups. Baboons were not fed except during capture. Two criteria were used to capture. First, a GPS weight limit of no more than 3% of body weight was adopted. As a result, 29 infants (46.03% of the AI group) were excluded from the study. Second, we excluded pregnant and lactating females and older individuals to avoid adverse health effects of capture and anesthesia. Individuals were chemically immobilized using ketamine (15 mg/kg) and fitted with GPS loggers (e-Obs Digital Telemetry, Gruenwald, Germany). The collars were fitted to five adult males, ten adult females, six adolescent males, and five juveniles (two males and three females). Adults attached the collars accounted for 62.5% of the total adult population. Because there is little sex difference in social development in individuals under four years of age [15], the age class ‘ juvenile’ in this study included both males and females. The collars were equipped with a breakaway mechanism (Advanced Telemetry Solutions, Isanti, MN, USA), which automatically released them at the end of the study.

We measured the positions of individual baboons using GPS during the day (6:00–18:00) for 30 days, from July 19 to August 17, 2019. Several collars were dropped along the way, reducing the number of tracked individuals. However, the priority of this study was to analyze the spatial positions of the largest number of individuals [38]; thus we analyzed 82,095 positions, at 1-minute intervals, of all 26 individuals, until we recorded the first failed GPS collar on July 24.

### Group spread

At each timestamp, the linear distance between two individuals with GPS in a group was calculated for all combinations; the maximum linear distance between two individuals at a single timestamp was considered the group spread at that time.

### Predation risk proxies

Theoretical studies have proposed several proxies to predict the predation risk at different spatial positions (Table 1, Fig 1). We used the unlimited domain of danger (UDOD) and minimum convex polygon (MCP) to assess the predation risk for individuals at the edge of the group, where they are most likely to encounter predators (Fig 1a, 1b). The extent to which an individual was surrounded by others was then analyzed using the limited domain of danger (LDOD), surroundedness, and dyadic distances (Fig 1c, 1d, 1e). The details of each proxy are presented below.

**Table 1.** Summary of advantages and weaknesses of each predation risk proxy, and suggested cases of when to use each depending on the predator’s preferences.

| Proxy | Predator’s preference | Advantages | Weaknesses |
| --- | --- | --- | --- |
| Unlimited domain of danger (UDOD) | Predators attack the edge of the group (one or several individuals) | Simple ‘edge_ vs. ‘central_ individual definition | Infinite areas are ecologically unrealistic<br>Possibly unrealistic measure of predation perception<br>Distances to and locations of neighbors are unknown |
| Minimum convex polygon (MCP) | Predators attack the strict boundary of the group | Strict ‘boundary_ vs. ‘inside_ definition | Distances and locations of neighbors are unknown |
| Limited domain of danger (LDOD) | Predators attack isolated individuals (less neighbors and more widely spaced to them) | Includes information on the predator’s range of attack | Difficulty in defining the radius<br>Possibly unrealistic measure of predation perception<br>Limited information on the number of neighbors and their closeness (distance) |
| Surrundedness | Predators attack isolated individuals (less surrounded by neighbors) | Includes information on the position and direction of all other group members<br>More fine-scale measure of spatial centrality (vs. UDOD and MCP) | Distance to neighbors unknown |
| Dyadic distances | Predators attack isolated individual (more widely spaced to neighbors) | Includes information on the distances to all other group members | Location (direction) of neighbors are unknown |

**Fig. 1.**
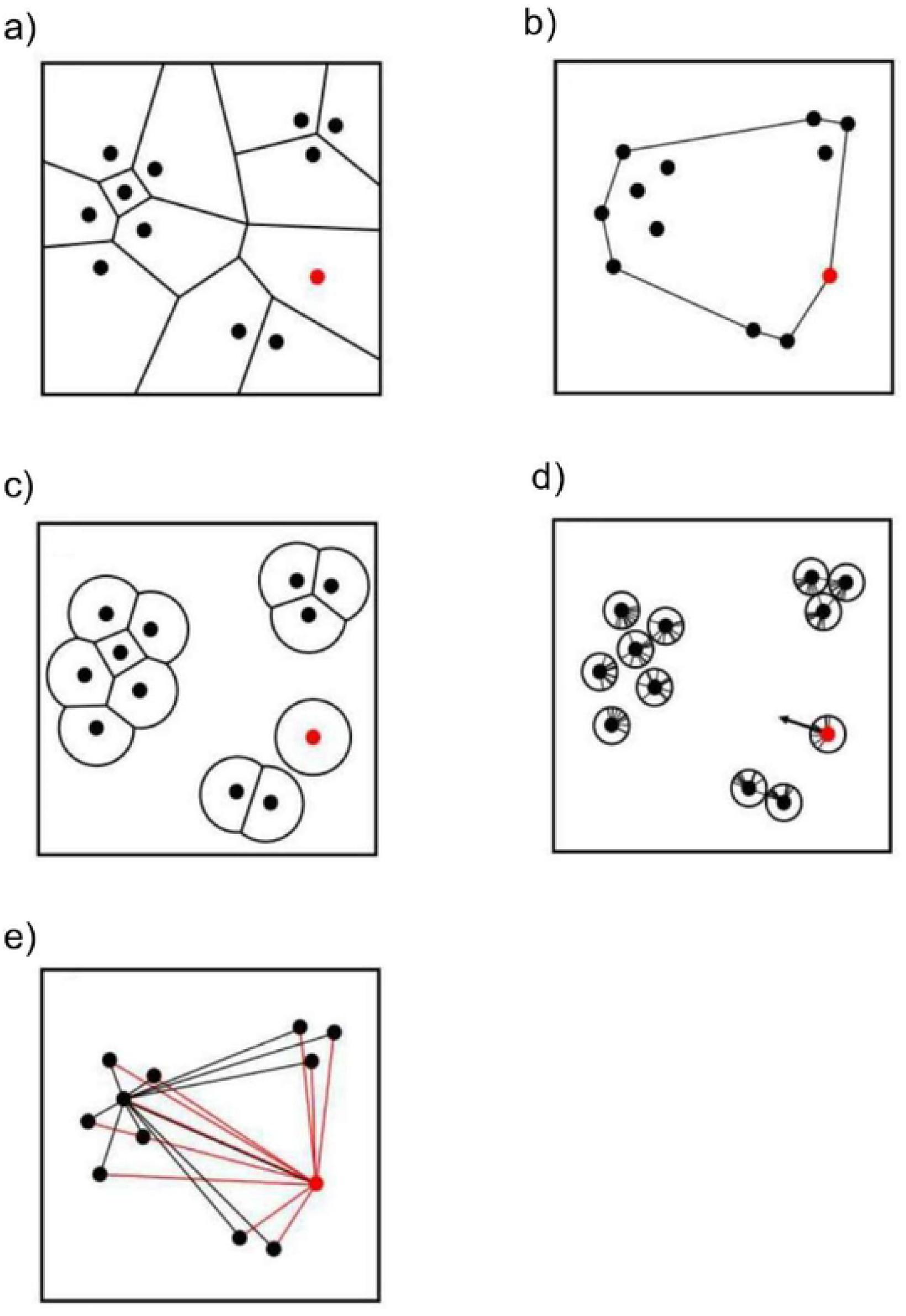
Schematic of each predatory risk proxy. The points represent individuals, a) unlimited domain of danger (UDOD); b) positions based on a minimum convex polygon (MCP); c) limited domain of danger (LDOD); d) surroundedness
(circular variance); e) dyadic distances. Points are individuals. If the red point is infinite in the UDOD or on the boundary in the MCP, it is the most vulnerable individual compared with its group mates Similarly, if the size of the LDOD is maximal, the surroundedness is high, or the dyadic distance is large (the black line is the dyadic distance), the individual in red is the most vulnerable compared to its group mates.

### a. Unlimited domain of danger (UDOD)

Hamilton [29] proposed the domain of danger (DOD) concept as a measure of the predation risk taken by each prey individual within a group. This concept assumes that a predator currently undetected by prey might appear anywhere within the field, inside or outside the group. DOD is defined as the space around the individual within which a predator can attack and kill it; the larger the size, the higher an individual’ s expected predation risk. The predatory is assumed to attack individuals at the closest distance (Table 1). When a perpendicular bisector is drawn on a line connecting points (mother points) placed at arbitrary neighboring positions, the nearest neighbor region formed around each mother point is called a Voronoi cell. When applying Hamilton’ s idea to real animal groups, a general problem arises: DOD is defined as Voronoi cells, but simultaneously, individuals at the edge of the group have biologically unrealistic unbounded Voronoi polygons (‘ unlimited domain of danger,’ UDOD) (Fig 1-a).

### b. Minimum convex polygon (MCP)

Predation risk is higher for individuals at the edge of a group than for those at the center [39]. One strategy to determine the individuals occupying these positions is based on a minimum convex polygon (MCP). In this method, animals were defined as being at the edge (boundary) of the group if they belonged to the set of vertices describing the smallest convex polygon enclosing all members of the group [40] (Fig 1b). Consequently, all the other individuals were defined as those inside the group.

### c. Limited domain of danger (LDOD)

Applying Hamilton’ s ideas to real animal groups raises the problem that the DODs of edge individuals are infinite, which is a biologically unrealistic assumption [41]. Therefore, subsequent theoretical studies have propounded an alternative proxy called the ‘ limited domain of danger’ (LDOD) [42]. The LDOD is the area of a circle of radius *r* centered on each animal. This radius corresponds to the range over which a predator can strike, the maximum detection distance of the predator, or the distance within which the predator can successfully launch an attack after approaching undetected [42]. If a focal individual is, its LDOD is maximal, defined as π*r*2 where *r* is the predator’ s attack distance (Fig 1c). If there is a conspecific individual within its maximum LDOD, the distance between the two is less than 2*r*, and the created bisector reduces the LDOD.

The definition of *r* based on field reports is important to represent the most realistic range of danger. Information on the attack ranges of baboons’ main ambush predators, leopards and lions, is scarce in the literature. However, only two studies have been cited. Observations by Bailey [43] suggest that prey will allow visible leopards to approach 25 m without feeling threatened but will flee if they are closer than 10 m. Specifically, there is a distance limit of 10–25 m within which a leopard can hide undetected by a baboon; therefore, 25 m was chosen for this study. Because anubis baboons also suffer from lion predation, we also calculated LDODs with a radius of 70 m based on the study by [44] on chacma baboons (*Papio ursinus*) and their use of refugia against lions and leopards. This value of 70 m is the median distance to a refuge, where chacma baboons living on open plains spend most of their time. This distance is hypothesized to be the distance beyond which a baboon is outrun and captured by a predator before reaching safety. Empirically, Stander [45] reported that 73% of lion kills had short chase distances of less than 20 m, whereas the remaining 27% were between 20–150 m. Moreover, Scheel [46] reported hunting distances of up to 200 m. In this study, individuals’ LDOD can theoretically vary between a minimum of 0 m^2^ (if at least two individuals share the same location) and a maximum of 1,960 and 15,369 m^2^ (the individual is isolated), with radii of 25 and 70 m, respectively.

### d. Surroundedness

‘ Surroundedness’ refers to the degree to which an individual is surrounded by all group members, with the expectation that more surrounded individuals should be less vulnerable to predation. This study assessed surroundedness using circular variance, a measure based on circular statistics [47].

The following calculations were performed according to Cremers and Klugkist [48]. A unit was drawn around a focal individual, and the directions of all the group members were projected as points on the circle’ s circumference. Connecting these points to the origin (the focal individual) yielded individual vectors (Fig 1-d). Vectors with the same value were stacked in the radial direction of the base circle. The resulting vector (R) was created by connecting the toe of the first vector to the head of the last vector. Relative to the focal individual, the direction indicated where most other group members belonged.

In this study, we calculated circumpolar variance, defined as 1-*R*. The length of the mean resultant vector (*R*) is R divided by the number of created vectors. A variance of 1 means that the individual is surrounded by many individuals, whereas a variance of 0 means that the individual is not surrounded; that is, all other members are located in one particular direction.

### e. Dyadic distances

Predation risk will likely vary with the number and proximity of conspecifics through effects such as dilution and accelerated predator detection [32]. Thus, one way to determine which individuals are more susceptible to predation is to determine their distance from all members. Individuals that maintain a greater distance from others are more physically isolated (Fig 1-e) and thus may be the preferred target for predators. Therefore, we calculated the Euclidean distance for each focal individual and the dyadic combination of all other group members at each timestamp.

## Statistical analyses

We aimed to explore the potential differences among age-sex classes for all predation risk proxies. Because LDODs and surroundedness are continuous variables, we used linear mixed models (LMMs). The autocorrelation structure of the measured sample points was analyzed, and the data were fitted using a Gaussian model with the following equation:

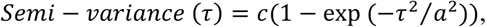

where c, a, and τ represent sill, range, and time-lag (min). These calculations were conducted at each timestamp (every 1 min interval) using the Python 3 (version 3.7.10) libraries NumPy (version 1.20.2), Pandas (version 1.3.5), and Matplotlib (version 3.3.4). We set these variables as the response variables, with age-sex classes (factors with three levels: adult males, adult females, adolescent males, and juveniles) as predictors. The interaction term between age and sex was also included. Because both the UDODs and MCP positions are binomial factors, we used generalized linear mixed models (GLMMs). Both models were fitted using a binomial error structure, with the response variable being the individual spatial position. The UDOD was defined as finite UDOD [0] vs. infinite UDOD [1], and the MCP positions were inside [0] vs. boundary [1]. The predictors were similar to those of the LMMs. To study the differences in dyadic distances between each age-sex class, we used an LMM with dyadic distances as the response variable and pairs of age-sex classes (factors with 10 levels) as the predictor. A two-sided test was used to ascertain the differences in mean values for each age-sex group. All LMMs and GLMMs were followed by Tukey’ s post-hoc comparisons with Bonferroni correction to assess the specific differences between each age-sex class.

In all the models, individual identities were set as random factors on the intercept. The significance of predictors in each model was computed with the model including the predictor against the model excluding the predictor (Wald χ2 for LMMs and likelihood-ratio χ2 tests for GLMMs, Anova Type III). Marginal R2 and conditional R2 were given for each model, representing the variance explained only by fixed effects and the variance explained by fixed and random effects, respectively. Results were considered significant at the α = .05 level for the models and post-hoc comparisons.

R packages (version 4.1.1; R Core Team [49]) used for the analyses were ‘ lme4’ for setting LMMs and GLMMs, ‘ multcomp’ for the post-hoc analyses, ‘ car’ for ANOVA tables and ‘ MuMin’ for the computation of R2.

## Results

### Group spread

The median group spread across all timestamps was 205.18 m (136.20 m in the first quartile and 345.18 m in the third quartile). The maximum and minimum distances were 3,560.79 and 20.89 m, respectively.

### Risk of exposure to predation

#### a. UDODs

Infinite UDODs were the most common among adult males, followed by adult females, adolescent males, and juveniles (Fig 2a, Supplementary Material S1). Significant differences were in the percentage of finite UDODs among the age-sex classes (x2 = 50.40, p < .001). The adult males had the smallest proportion of finite UDODs (vs. adult females, z = 4.63, p < .001, vs. adolescent males, z = 5.87, p < .001, vs. juveniles, z = 6.33, p < .001). No significant differences were observed in any other comparisons.

**Fig. 2.**
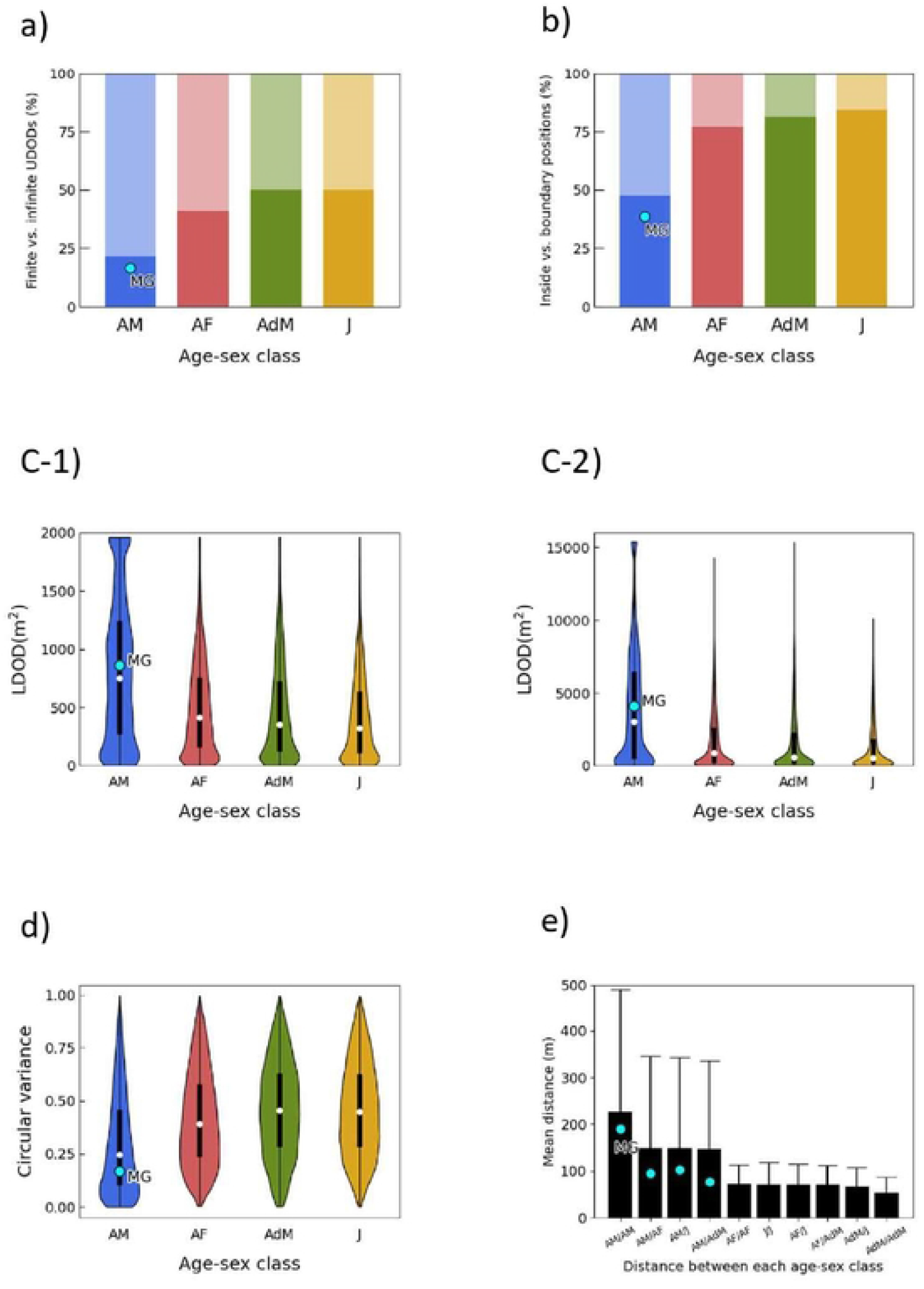
a) Percentages of finite (dark gray) vs. infinite (light gray) UDODs for each age-sex class, ordered from lowest to highest Age-sex class abbreviations are as fol lows: AM = adult males, A F = adult females, AdM = adolescent males, and J = juveniles. N oration is similar in subsequent figures, b) Percentages of the inside (dark gray) vs. boundary (light gray) MCP positions for each age-sex class, c-1, c-2) Violin plots along boxplots of LDODs. c-1) radius of 25 mand c-2) radius of 70 m ⇃ denotes the mean value, d) Violin plots along box plots of surroundedness. e) Mean distance (m) between each age-sex class Bars are mean associated with their standard error.

#### b. MCP

Significant differences were in the percentage of individuals in the edge/center among the age-sex classes (x2 = 83.52, p < .001, Fig 2b, S2). Adult males were more frequently on the edge than individuals of other age-sex classes (vs. adult females, z=7.36, p < .001, vs. adolescent males, z=6.86, p < .001, vs. juveniles, z=8.04, p < .001). No significant differences were observed in any other comparisons.

#### c. LDODs

For both LDODs with radii of 25 m (attack of a leopard) and 70 m (attack of a lion), there were differences among the age-sex classes (Fig 2c1, 2c2). For LDODs with a radius of 25 m, significant differences were in the percentages of finite LDODs among the four age-sex groups (x2 = 606.76, p < .001, S3). Adult males had the largest LDODs (vs. adult females, z = 18.67, p < .001, vs. adolescent males, z = 19.75, p < .001, vs. juveniles, z = 22.36, p < .001). Similar qualitative differences were found for the 70 m radius LDODs (x2 = 41.93, p < .001, vs. adult females, z = 4.94, p < .001, vs. adolescent males, z = 5.22, p < .001, vs. juveniles, z = 3.91, p < .001). No differences were found at either 25 m or 70 m in any of the other comparisons.

#### d. Surroundedness

There were significant differences in the degree of being surrounded by group members among the age-sex classes (x2 = 343.16, p < .001, Fig 2d, S4). Adult males had the smallest circle variances (vs. adult females, t=13.86, p < .001, vs. adolescent males, t=-16.16, p < .001, vs. juveniles, t=-15.80, p < .001). No significant differences were observed in any other comparisons.

#### e. Dyadic distances

Of all the age-sex classes, the mean distance between adult males was the greatest, and between adolescent males was the smallest (Fig 2f). Distances between adult males (mean ± SD, 226.38 ± 526.76 m) were significantly longer than that between adult females (71.45 ± 84.02 m, z=3.31, p < .005, S5) or that between adolescent males (53.66 ± 67.96, z=3.30, p < .005). Statistical differences were not found between juveniles (70.83 ± 93.02, z=3.31, p < .005).

### The special position of the alpha male

The UDOD and MCP results showed that the alpha male, *MG*, was located peripherally of the group to a similar extent as the other males (Fig 2a, 2b). In addition, LDOD, surroundedness, and dyadic distance indicated that the *MG* was not surrounded less by individuals as other males.

## Discussion

In Hypothesis 1, we proposed that adult males are located peripherally, where coursing predation is most likely to occur. Our results from the UDOD and MCP showed that, as expected, adult males were located more on the periphery, consistent with previous studies on primates [50] and other animals [51–53]. Two well-studied hypotheses have been proposed from ecological and sociological perspectives to explain why males are located peripherally. One hypothesis is that there is a trade-off between increased foraging efficiency and higher predation risk [52,54]. Adult males with larger bodies and higher nutritional requirements may have prioritized greater food accessibility over predation risk and occupied peripheral positions. In addition, adult males may play a role in ‘ protecting’ females and their young in the group’ s center through vigilance, alarm, mobbing, and counterattacks against predators or neighboring groups [55]. Four reasons have been cited for this behavior: paternal investment, maintenance of males and female social bonds, group enhancement benefits, and quality signaling in the mating market [56].

Hypothesis 2 stated that adult males were the most isolated individuals vulnerable to ambush predators. As expected, the LDOD, surroundness, and dyadic distance indicated that adult male baboons were the most isolated. One reason for this may be the social structure of baboons. In all four species of savanna baboon, anubis, yellow, chacma, and guinea baboons, females remain in their natal groups throughout their lives, whereas males generally move to another group after sexual maturity. In species with male dispersal, adult males are expected to have few or no relatives in the postnatal group except for their offspring. DNA microsatellite association analyses in some primates have reported that adult males have few close relatives in the postnatal groups [57,58]. As unrelated males are severe competitors of females in estrus, adult males maintain a long distance from each other to escape attacks by other males and serious injuries [59].

Hypothesis 3 states that individuals with high social status among adult males, especially alpha males, will occupy the secure core of the population. Dominant individuals, who can access preferential foods even under intense competition, are expected to be in the safety center of the group [31,60–62]. However, the alpha male in this study was peripheral and isolated, as were other males. Intimacy in the male-female relationship, called ‘ friendship’ in baboons [63], prevents male isolation, and this relationship in yellow baboons is formed through the offspring [64]. One possible explanation for our result may be that this alpha male, although born in this group, had been in a position for only approximately one year and had few female friends and offspring.

We addressed the methodological and theoretical advantages and disadvantages of the proxies used in this study for each hunting style of the predators. A strong assumption is that there is variation in predation risk based only on spatial positioning, for which predators can show variable preferences. First, assuming a coursing predator, UDOD and MCP positions are similar in that they use a simple dichotomy to define individuals located on the edge of the group as opposed to those centrally positioned. However, both methods vary in how they define edges owing to their underlying mathematics, which can result in slightly different results. For example, Voronoi diagrams can define more individuals as peripheral (‘ edge’ individuals) compared to MCP vertices that only consider peripheral individuals as those strictly forming the spatial boundary of the group; individuals with finite UDODs are always inside the group. These differences in definition are illustrated by the left and top-right clusters of individuals in Figure 1a and 1b, respectively. Using one proxy will then depend on whether the predator prefers to strictly attack the individual forming the group’ s boundary or any individual forming the periphery (‘ edge’), including the one at the strict boundary. Both methods are useful only when predators consistently attack the edge of a group (marginal predation). Indeed, this has been observed in diverse species [29,65–67].

From a theoretical perspective, LDODs are based on the notion that animals react to the open spaces around them, whereas the surroundedness and dyadic distance highlight the possible effects of several neighbors. In a context where predators do not consistently attack peripheral individuals but choose isolated individuals, those outside or inside the group, who are relatively distant from group members, other variables may be more pertinent. LDODs can be useful because they include information on the distance to neighbors regardless of the position of the individual within the group. However, its calculation can be complicated if the group has several types of predators with different hunting techniques. Consequently, it raises the question of whether one should expect changes in results when comparing groups of different radii. As such, the radius will rest on a detailed description of predator-prey interactions and will be somewhat arbitrary.

Considering that LDODs only include information on the closest neighbors, the surroundedness and dyadic distances complement each other and provide a better measure of isolation based on the positions of group members. Surroundedness is a more fine-scale measure of spatial centrality/peripherality within a group (especially compared to UDODs and MCP positions) and includes information on the position and direction of all other group members. Dyadic distances are a more fine-scale measure of neighbor spacing and include information on the distance to all other group members. In some species, isolation, rather than the periphery, is correlated with higher predation risk, such as in straggler fish, that is, isolated individuals outside the school, in which centrally positioned members of the school are also more at risk [23], or in redshanks, where preferred targets for predators are those that are more widely spaced compared to their nearest neighbors [68].

The number of collared individuals is the most limiting factor in emphasizing these predation risk proxies and any other proxies correlated with predation risk. Missing individuals (non-collared) influence the shape of the Voronoi diagrams and MCPs, change the size of the LDODs, and modify the circular variance and distances between individuals. Because predation risk is always related to all other group members, it is necessary to know the location of the maximum number of, if not all, individuals to be as accurate as possible. However, this study prioritized the maximum number of individuals, which has the disadvantage of reducing the number of data days. Farine et al. [30], who conducted a similar data collection and analysis, noted that individuals showed a consistent pattern of spatial positioning within the population over many days. However, in the present study, the maximum h-spread of the group was slightly greater than 3 km, which was attributed to an older male lagging behind the group by approximately half a day. Excluding this day’ s data, the median and maximum spread distances for the group were 184.83 and 734.11 m, respectively. Because reducing the number of days of analysis by a factor of h can cause such variation, it is necessary to develop a methodology to determine the time interval of positioning suitable for spatial location analysis and the number of valid data days.

Elucidating the costs of predation risk associated with individuals’ spatial positions within a group is important for understanding group formation and maintenance. In this study, we demonstrated that adult males were consistently weaker than predators that attacked them from outside the group, as well as predators that ambushed them within the group. Determining whether adult males position themselves voluntarily or unwillingly on the edge is difficult; however, it is an interesting future challenge for both ecological and social interests.

## Supporting information

S1 Percentages of infinite positions are given for UDODs and results of GLMM.

S2 Percentages of boundary positions and results of GLMM.

S3 Summary statistics of LDOD size and results of LMM.

S4 Summary statistics of Surroundedness and results of LMM.

S5 Mean distances (m) of each pair of age-sex classes.

S6 Data and codes for analysis (Python and R) (ZIP).

## Author Contributions

Conceptualization: Alexandre Suire, Akiko Matsumoto-Oda Data curation: Roi Harel, Margaret Crofoot

Formal analysis: Alexandre Suire, Itsuki Kunita

Founding acquistition: Margaret Crofoot, Akiko Matsumoto-Oda

Investigation: Roi Harel, Mathew Mutinda, Maureen Kamau, James M. Hassel, Suzan Murray, Akiko Matsumoto-Oda Methodology: Alexandre Suire, Itsuki Kunita, Akiko Matsumoto-Oda

Project administration: Roi Harel, Margaret Crofoot, Akiko Matsumoto-Oda Supervision: Akiko Matsumoto-Oda

Validation: Alexandre Suire, Itsuki Kunita, Roi Harel, Margaret Crofoot, Mathew Mutinda, Maureen Kamau, James M. Hassel, Suzan Murray, Akiko Matsumoto-Oda

Visualization: Alexandre Suire, Itsuki Kunita, Akiko Matsumoto-Oda Writing – original draft: Alexandre Suire, Akiko Matsumoto-Oda

Writing – review & editing: Alexandre Suire, Itsuki Kunita, Roi Harel, Margaret Crofoot, Mathew Mutinda, Maureen Kamau, James M. Hassel, Suzan Murray, Akiko Matsumoto-Oda

## Acknowledgments

We thank National Commission for Science, Technology and Innovation (Kenya), Kenyan Wildlife Service, Institute of Primate Research (MNK, Kenya), and Mpala Research Centre for permission to conduct research. We also thank Carter Loftus, Shauhin Alavi, Agrey Masiva, Rebecca Stites, Ellie Milnes, Jenn Yu, Sharon Mulindi, Nashipai Seketeti, Peter Ombewa, Andrea Surmat, Lenareiyo Ltukulen and Zakayo Parsimey for their assistance in the baboon captures. We also thank Reiji Suzuki and Eiiti Kasuya for advice in the analysis.

